# Temporal Trajectories of Motor and Cognitive Dysfunction After Combined PTSD and TBI in Mice: Implications for Neurodegenerative Disease Vulnerability

**DOI:** 10.64898/2026.06.03.729686

**Authors:** Kayla L. McDaniel, Carolyn E. Tinsley, Laura Dovek, Zoe Potter, Lorenzo R. Nungaray, Nathan M. McGuire, Jasmine Loeung, Peyton T. Wickham, Jonathan E. Elliott, Charles K. Meshul, Miranda M. Lim

## Abstract

Post-Traumatic Stress Disorder (PTSD) commonly occurs alongside Traumatic Brain Injury (TBI), yet the chronic behavioral consequences of combined neurotrauma (i.e., PTSD and TBI) across the sexes remain unclear. Using a mouse model combining Single Prolonged Stress (SPS) as a model for PTSD and Controlled Cortical Impact (CCI) as a model for TBI, we assessed gait, anxiety-like behavior, and contextual fear learning and extinction at 2, 4, and 12-weeks post-injury. Combined neurotrauma produced early and persistent gait impairments in both sexes, delayed changes to anxiety-like behavior characterized by reduced avoidance of an anxiogenic environment, and long-lasting contextual fear recall deficits. Impaired learning was observed in males, where they demonstrated reduced fear acquisition and diminished extinction rates at later time points while females showed no deficits. Across testing and sex, peak deficits emerged at 4 weeks post-neurotrauma. Together, these findings define a sex- and time-dependent behavioral phenotype following combined neurotrauma and underscore the importance of modeling comorbidity to capture the temporal and neurobehavioral consequences of trauma exposure that more closely reflect clinical populations.

## Introduction

Post-traumatic stress disorder (PTSD) is characterized by persistent disturbances in fear regulation, emotional processing, and various cognitive and memory impairments that, by definition, require a period of post-trauma incubation. A large body of clinical literature exists describing the delayed and prolonged impact of PTSD on neurocognitive functioning in trauma-exposed populations.^1-4^ In rodent models of PTSD, alterations in cognitive ability, fear responding, and avoidance behavior often emerge or become most reliably detectable after an incubation period.^5-8^ Sex differences are also prominent. Females are more likely to meet criteria for PTSD after trauma exposure than males^9-12^ and preclinical studies demonstrate sex-dependent differences in fear expression, extinction retention, and stress physiology.^13-15^

In clinical populations, Traumatic brain injury (TBI) is increasingly recognized as a chronic condition with long-lasting behavioral consequences that extend beyond the acute post-injury period. Beyond cognitive symptoms, TBI is associated with persistent behavioral and neuropsychiatric disturbances that substantially affect long-term recovery and quality of life.^16-23^ Importantly, these outcomes often evolve over weeks to months and even years, highlighting the importance of considering the elapsed time post-injury when assessing behavioral variables.^24-26^ Sex also represents a critical biological variable in TBI outcomes with males twice as likely to sustain TBI requiring hospitalization than females (CDC^27^). Preclinical studies report sex-dependent differences in neuroendocrine and neuroinflammatory responses to injury,^28-32^ and clinical studies similarly suggests differences in symptom burden and recovery trajectories between males and females.^33-36^

Critically, TBI and PTSD frequently co-occur in clinical populations, particularly among military service members and Veterans, where comorbidity is common and associated with greater symptom severity, poorer functional outcomes, and increased long-term disability compared with either condition alone.^4, 20, 37-41^ This comorbid presentation represents one of the most prevalent and understudied trauma-related clinical conditions, yet remains poorly modeled in preclinical research. ^42-45^

Preclinical models of combined TBI and PTSD-like trauma have provided an important framework for examining how these exposures interact to produce persistent behavioral and neurobiological dysfunction. In an early study, Ojo et al. developed a mouse model combining repeated predator-odor exposure, restraint, and inescapable foot shock with concussive impact TBI, and reported increased anxiety-like behavior and neuroinflammatory markers following combined exposure.^43^ Subsequent preclinical models have used different combinations of stress and injury in mice and rats, including single prolonged stress (SPS) with controlled cortical impact (CCI),^45^ chronic variable stress with closed-head injury^44^ and SPS with closed-head weight-drop injury.^42^ Across these approaches, combined neurotrauma generally produces behavioral and biological impairments that are greater than, or distinct from, those observed after either PTSD or TBI alone. However, most prior studies have been conducted exclusively in males, and experimental designs vary substantially in the timing, sequence, severity, and post-injury interval used to model combined trauma, limiting direct comparison across studies.

The present study extends this model by systematically examining sex- and time-dependent behavioral trajectories following the combined TBI and PTSD model described by Teutsch et al 2018 (SPS+CCI).^45^ By assessing gait, anxiety-like behavior, and contextual fear acquisition and extinction at 2, 4, and 12-weeks post-neurotrauma in both male and female mice, we aim to define the temporal emergence of behavioral dysfunction after combined neurotrauma.

## Methods

### Subjects

Male and female C57BL6/J mice were used in all experiments. Mice were partially sourced from Jackson Laboratory and partially sourced from our in-house breeding colony. All outcomes presented here were examined for differences in sourcing and none were found. Mice were between 60-120 days old at the start of the experiment. Age was tracked as a covariate and when significant is reported below. Housing was in a 12h:12h light:dark cycle (lights on at 0700 h) with ad libitum access to food (LabDiet PicoLab Laboratory Rodent Diet 5L0D) and water. Temperature (21.6C°) and humidity (∼24%) were monitored and maintained at constant levels. All experiments and housing conditions were approved by the Institutional Animal Care and Use Committee of the VA Portland Health Care System and adhered to guidelines set forth by the National Institutes of Health Guide for the Care and Use of Laboratory Animals. N=182 animals started the study.

### Experimental Design

At the beginning of the experiment, mice were split (approximately equally) into neurotrauma or control conditions with attempts to counterbalance source, breeder pairs, age, and sex. The neurotrauma group underwent a combined neurotrauma model of PTSD and TBI – single prolonged stress (SPS) and controlled cortical impact (CCI), respectively (details below). Due to the methodological complexity and swimming component of SPS, the SPS protocol was performed first, followed by CCI.

Following SPS+CCI, subjects were left undisturbed except for weekly cage changes in standard home cages for either 2, 4, or 12-weeks depending on their group. After the assigned period of time passed, mice underwent behavioral tests: Light/Dark Box, DigiGait, and Contextual Fear Conditioning and extinction (see **Fig. 1** for experimental design). Each test was separated by approximately 24 hours and was conducted during the light phase. At the conclusion of the experiment, mice were euthanized by an overdose of inhaled isoflurane and tissue collected for future analysis.

**Figure 1:**
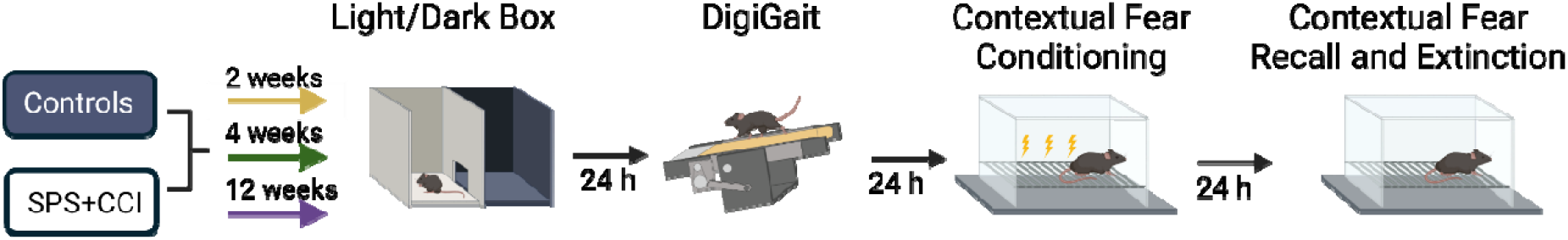
Experimental Design. Mice underwent single prolonged stress (SPS) followed by controlled cortical impact (CCI) to model PTSD+TBI, respectively. A behavioral battery to examine anxiety like behavior (Light/Dark Box), gait (DigiGait), and cognition (contextual fear conditioning followed by extinction) was conducted either 2, 4, or 12 weeks after CCI. Each cohort was run with sham Controls that followed identical timing. Schematic prepared with Biorender.

### Single Prolonged Stress (SPS)

SPS was used as a model of PTSD.^8, 46-50^ SPS consists of four sequential stressors: 2-hour tube restraint, group forced swim, ether anesthesia until loss of consciousness, and social isolation. During SPS, Control mice were transported with the experimental group but were not exposed to the SPS model. For tube restraint stress, mice were placed into ventilated 50mL conical vials, which were then placed in a clean cage with bedding, for two hours. After tube restraint, groups of 4-6 mice were immediately placed together into a single plastic tub (8.5 × 9.0 × 12.0 in) filled with room temperature water (26°C), deep enough to prevent tails from touching the bottom (three-quarters full). Group forced swim lasted a duration of 15 minutes, during which time the mice and water temperature were closely monitored. Following the group forced swim, mice were towel dried and allowed to rest for ∼15-20 minutes. Finally, mice were individually secured in a jar and exposed to 1.0mL of diethyl ether until loss of consciousness. Mice were carefully monitored and immediately removed from the jar once they lost consciousness. Subsequently, mice were placed in a standard home cage and individually housed.

### Controlled Cortical Impact (CCI)

A moderate TBI was induced approximately 3-5 days after SPS with controlled cortical impact (CCI). Mice were weighed and anesthetized with isoflurane (3-5% induction, 1-3% maintenance) in oxygen (1 L/min). Once anesthetized, mice were secured on a stereotactic frame and optical lubricant was applied.

The hair on the animal’s head was shaved, and the skin was sterilized using 70% alcohol followed by iodine in triplicate. After cleaning, topical lidocaine/prilocaine cream was applied to the scalp. Once there was a clear and sterilized window on the top of the animal’s head, a small incision was made, revealing the skull. A single use cotton swab with 0.3% peroxide was applied to the skull to remove membranes, revealing bregma and lambda. A hand drill was used to create a small craniotomy in the bottom right quadrant. An impactor arm (Kopf) was aligned with the craniotomy and used to deliver an impact at a depth of 2mm with a velocity of 4.5m/s and a dwell time of 0.1s. Following impact, skulls were cleaned and dried using single use cotton swabs before suturing. Once sutures were in place, a small amount of super glue was placed over the sutures to ensure successful wound closure. The mice were given topical lidocaine and antibiotic ointment before returning to a sanitized home cage. The mice were given 1ml of children’s cherry flavored acetaminophen (Q-Pap) in their drinking water daily for up to 7 days. Mice in the Control group were anesthetized under isoflurane for 10-15 minutes, but they did not undergo a surgical procedure. N=3 animals were removed from the experiment and euthanized as a result of complications from the CCI procedure or recovery.

### Timing Post-Neurotrauma

Following CCI, mice were assigned a timed incubation period before receiving behavioral testing. Mice were counterbalanced to either a 2-, 4-, or 12-week time point. Mice in the SPS+CCI group were single-housed for the duration of the experiment, and the Control groups were group-housed in cages of 2-4.

### Light/Dark Box

Anxiety-like behavior was measured by a 10 minute test inside a Light/Dark (LD) Box consisting of an open-field chamber (60 cm x 60 cm x 30 cm; Omnitech Electronics) partitioned with a darkened area consisting of a black insert (30 cm x 30 cm x 20 cm). Infrared beams on all sides measured locomotor activity and location. A cutout door (10 cm x 5 cm) allowed the mouse to move freely between zones. Mice were placed in the light portion facing the wall furthest from the door where time and activity were measured for each zone (light or dark). All mice were habituated to the testing room for 30 minutes prior to testing. n=13 animals were excluded due to software malfunctions but were still exposed to the testing apparatus for the same amount of time. Subjects used in analysis include the following:

2-w females: n=16

2-w males: n=17

4-w females: n=6

4-w males: n=14

12-w females: n=9

12-w males: n=12

Control females: n=38

Control males: n=54

### Gait Testing

24 hours after LD box testing, each subject underwent gait testing using the DigiGait™ imaging system (Mouse Specifics). The DigiGait system consisted of a transparent treadmill belt surrounded by a plexiglass compartment (17.4 cm× 5.1 cm× 13.3 cm) that was illuminated from below. The treadmill is set to a 15-degree decline and a speed of 25 cm/s. For this test, mice were placed in the compartment and allowed to habituate for 1 minute. After the habituation period, the treadmill was turned on, and each animal was recorded for approximately 5 seconds. N=74 animals were excluded due to a computer failure that corrupted data, all animals were exposed to the gait apparatus for the same amount of time. Gait was analyzed using the DigiGait Analysis software (version 16) according to manufacturer’s instructions by a trained researcher blind to group and sex. Subjects used in analysis include the following:

2-w females: n=7

2-w males: n=7

4-w females: n=6

4-w males: n=9

12-w females: n=8

12-w males: n=11

Control females: n=27

Control males: n=30

### Contextual Fear Conditioning and Extinction

The contextual fear conditioning and extinction paradigm occurred over two days. Contextual Fear Acquisition: On the first day of fear conditioning, mice were brought into the testing room to habituate to the room for 30 minutes before the start of the experiment. Following room habituation, mice were weighed and then placed individually into the testing chambers. Four identical testing chambers (24 cm x 30 cm x 21 cm; MedAssociates) were used for fear conditioning which consisted of two clear acrylic walls, two stainless steel walls, and a clear top. Chambers were enclosed by sound attenuating cubicles (MedAssociates). Chambers were lit by a white house light and equipped with fans that provided background white noise of 65 dB. Each chamber had a metal floor that consisted of 36 stainless steel rods spaced approximately 5 mm apart connected to a shock generator. The unconditioned stimulus (US) was a 1.0 mA scrambled footshock (duration = 1s). Mice were placed in chambers and after 3 minutes received n=5 US deliveries with a fixed intertrial interval (ITI) of 60s. Mice were removed exactly 60 seconds after the termination of the last shock and returned to their home cages. Chambers were cleaned with Nolvasan solution between each trial to remove any residual scent cues.

Fear recall and extinction: On the second day of fear conditioning, mice were brought into the testing room and allowed to habituate for 30 minutes prior to testing. After room habituation, each mouse was placed back into the conditioning chamber for a 20-minute extinction session. No shocks or other stimuli were presented during this period. Chambers were cleaned with Nolvasan solution between each trial to remove any residual scent cues. After testing, mice were returned to their home cages.

Freezing was scored manually by trained researchers blind to group and sex and was defined as the cessation of movement with the exception of respiration. N=29 animals were excluded due to hardware or software malfunctions. Subjects used for analysis included the following:

2-w females: n=14

2-w males: n=14

4-w females: n=9

4-w males: n=13

12-w females: n=6

12-w males: n=10

Control females: n=36

Control males: n=48

## Results

### Light Dark Box

The Light-Dark box was used to measure anxiety like behaviors in our different groups and was analyzed using ANOVA. When analyzing amount of time spent in the dark region of the light dark box, mice spent significantly less time in the dark chamber of the LD box 12 weeks after combined SPS+CCI than Controls (main effect of group: *F*(3,158)=3.38, *p*=0.020; post hoc Dunnett’s test compared to Controls: 2 weeks: *p*=0.453, 4 weeks: *p*=0.851, 12 weeks: *p*=0.005) (**Fig. 2a**). There were no effects of sex (main effect of sex: *F*(1,158)=0.06, *p*=0.814) nor a significant sex by group interaction (sex x group interaction: *F*(3,158)=2.06, *p*=0.108) of entries into the light chamber and observed no significant effects of group (main effect of group: *F*(3,158)=0.60, *p*=0.61), sex (main effect of sex: *F*(1,158)=1.71, *p*=0.193), or interaction (sex x group interaction: *F*(3,158)=0.22, *p*=0.884) on the number of entries into the light chamber made by mice (**Fig. 2b**).

**Figure 2:**
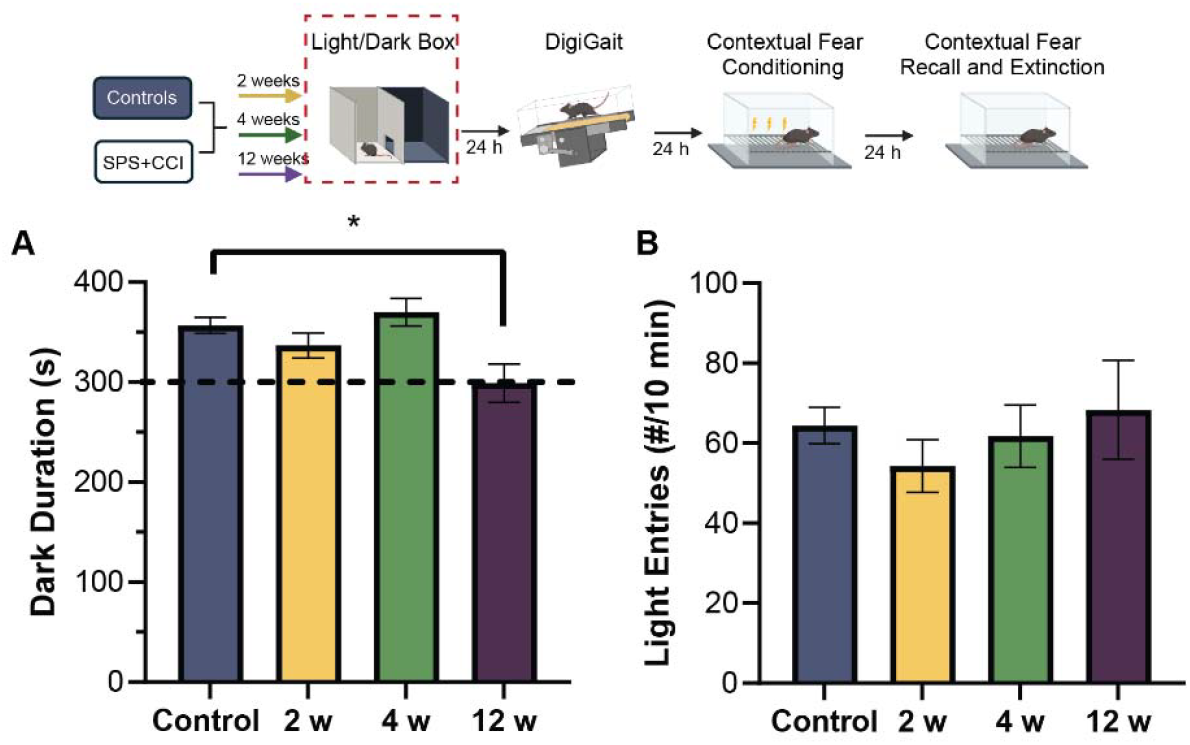
Delayed alterations in anxiety-like behavior following combined neurotrauma. A) Time spent in the dark compartment of the LD box. SPS+CCI exposed mice showed a significant reduction in dark duration at 12 weeks compared to Controls indicating altered responsivity to the anxiogenic environment. B) Number of light compartment entries during the LD box test did not differ between groups suggesting that changes in dark preference were not driven by changes in locomotor activity. No sex differences were observed. Data presented as mean +/-SEM. *p<0.05. Schematic prepared with Biorender.

### Impaired Gait After Neurotrauma

Primary gait metrics of interest were chosen *a priori* based on previous work on this topic (Teutsch et al 2018^45^): Paw placement positioning (PPP) and stance width. Neither of these metrics were influenced by age in Control mice. PPP was analyzed with ANOVA (between group factors = sex, trauma group) on each side. There was a main effect of trauma group (main effect of trauma: *F*(3,97)=17.36, *p*<0.001) on PPP on the left side (the side contralateral to the CCI) with post-hoc tests revealing that all time points showed significantly *lower* PPP compared to Controls (Dunnett’s, all *p* values <0.003)(**Fig. 3a**). There was also a main effect of trauma group on the right side (main effect of trauma: *F*(3,97)=3.45, *p*=0.020) with mice in the 2 week time point showing significantly *higher* PPP than Controls (*p*=0.020) but not in the 4 week (*p*=0.499) or 12 week (*p*=0.105) groups, suggestive of minor compensatory mechanism. There were no effects of sex on either side (main effect of sex: Left: *F*(1,97)=0.12, *p*=0.731; Right: *F*(1,97)=0.04, *p*=0.836) and no sex x group interactions on either side (sex x group interaction: Left: *F*(3,97)=1.23, *p*=0.303; Right: *F*(3,97)=2.10, *p*=0.106).

**Figure 3:**
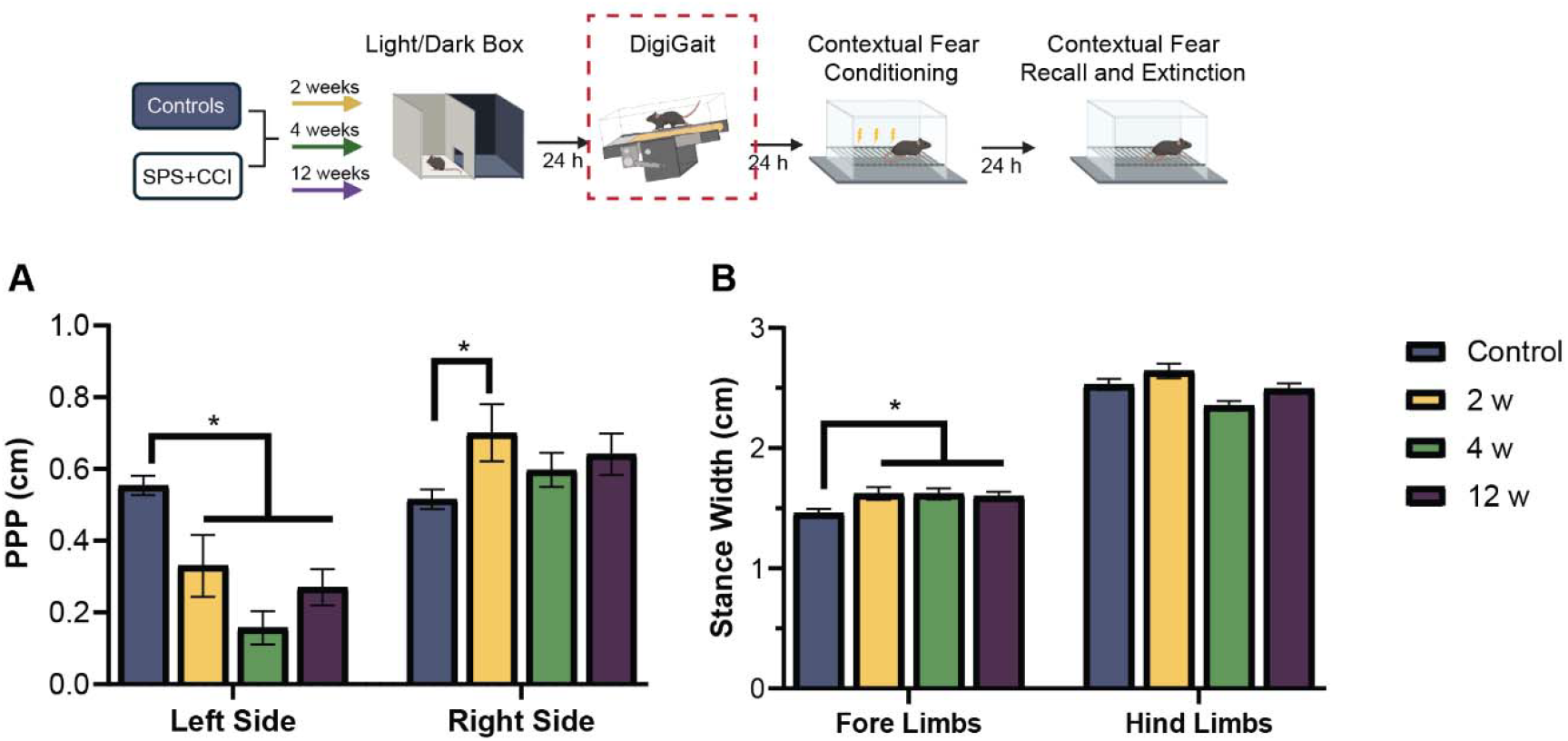
Impaired gait and balance metrics emerge at 2 weeks and persist until 12 weeks post SPS+CCI. A) Paw Placement Positioning (PPP) was decreased in the left side (side contralateral to CCI) at 2, 4 and 12 weeks after combined SPS+CCI compared to sham Controls and transiently increased at 2 weeks on the right side. B) wider stance width in the fore paws at all three time points after injury suggest postural adjustments for stability. Schematic prepared with Biorender.

Stance width was analyzed with separate ANOVAs (between group factors = sex, trauma group) on forelimb and hindlimb stance width. There was also a significant main effect of trauma group on fore limb stance width (main effect of trauma: *F*(3,97)=4.53, *p*=0.005), with all three time points showing significantly higher fore stance widths than Controls (Dunnett’s, all *p* values <0.039) (**Fig. 3b**). There was a trend towards a significant effect of sex (main effect of sex: *F*(1,97)=3.01, *p*=0.086) but no sex x group interaction (sex x group interaction: *F*(3,97)=0.83, *p*=0.480). In the hind legs, there was a trend towards an effect of trauma (main effect of trauma: *F*(3,97)=2.37, *p*=0.075) but similarly no effects of sex: (main effect of sex: *F*(1,97)=1.02, *p*=0.316; sex x group interaction: *F*(3,97)=1.12, *p*=0.346). Given the downhill orientation of the DigiGait treadmill, it is unsurprising that we see effects in the forelimbs but not the hind limbs.

### Impaired Contextual Fear Learning and Recall After Neurotrauma FC Day 1 Freezing

Freezing was measured for the entire duration of fear acquisition and percent freezing during each ITI (including the period prior to the first US and after the last US) and analyzed with repeated measure ANOVA (within subjects factor =US number; between subjects factor = neurotrauma group, sex). Assumptions for sphericity were not met, as such, *df* were Greenhouse-Geisser adjusted.

There was a significant within subjects effect of US number, with animals showing increasing levels of freezing at subsequent deliveries of the foot-shock (within subjects effect of US number: *F*(4.33, 619.28)=339.85, *p*<0.001) with no significant interaction of neurotrauma group or sex (within subjects US number x group interaction: *F*(12.99, 619.28)=1.32, *p*=0.194; within subjects US number x sex interaction: *F*(4.33, 619.28)=1.29, *p*=0.269). However, there was a significant US number x sex x neurotrauma group interaction (*F*(12.99, 619.28)=1.81, *p*=0.038) with males exposed to neurotrauma showing slower contextual fear learning during initial acquisition. Over the full conditioning session, there was a significant effect of neurotrauma group (between subjects, main effect of trauma: *F*(3,143)=3.28, *p*=0.023), no main effect of sex (between subjects, main effect of sex, *F*(1,143)=0.19, *p*=0.668), however there was a significant sex x neurotrauma group interaction (between subjects, sex x neurotrauma interaction: *F*(3,143)=2.80, *p*=0.042). Because of this significant group x sex interaction, post hoc testing (Dunnett, 2-sided) was performed on males and females separately. We found that across the entire contextual fear acquisition session males exposed to neurotrauma froze less than Controls 4 weeks after SPS+CCI (*p*=0.007) and showed a trend of decreased freezing compared to Controls after 12 weeks (*p*=0.070) but not at 2 weeks (*p*=0.912) (**Fig. 4a**). Interestingly, we found that females had heightened freezing during acquisition 2 weeks after neurotrauma compared to sex matched Controls (*p*=0.040, Dunnett). However their freezing returned to Control levels by week 4 (*p*=1.000), persisting into week 12 (*p*=1.000) (**Fig. 4b**).

**Figure 4:**
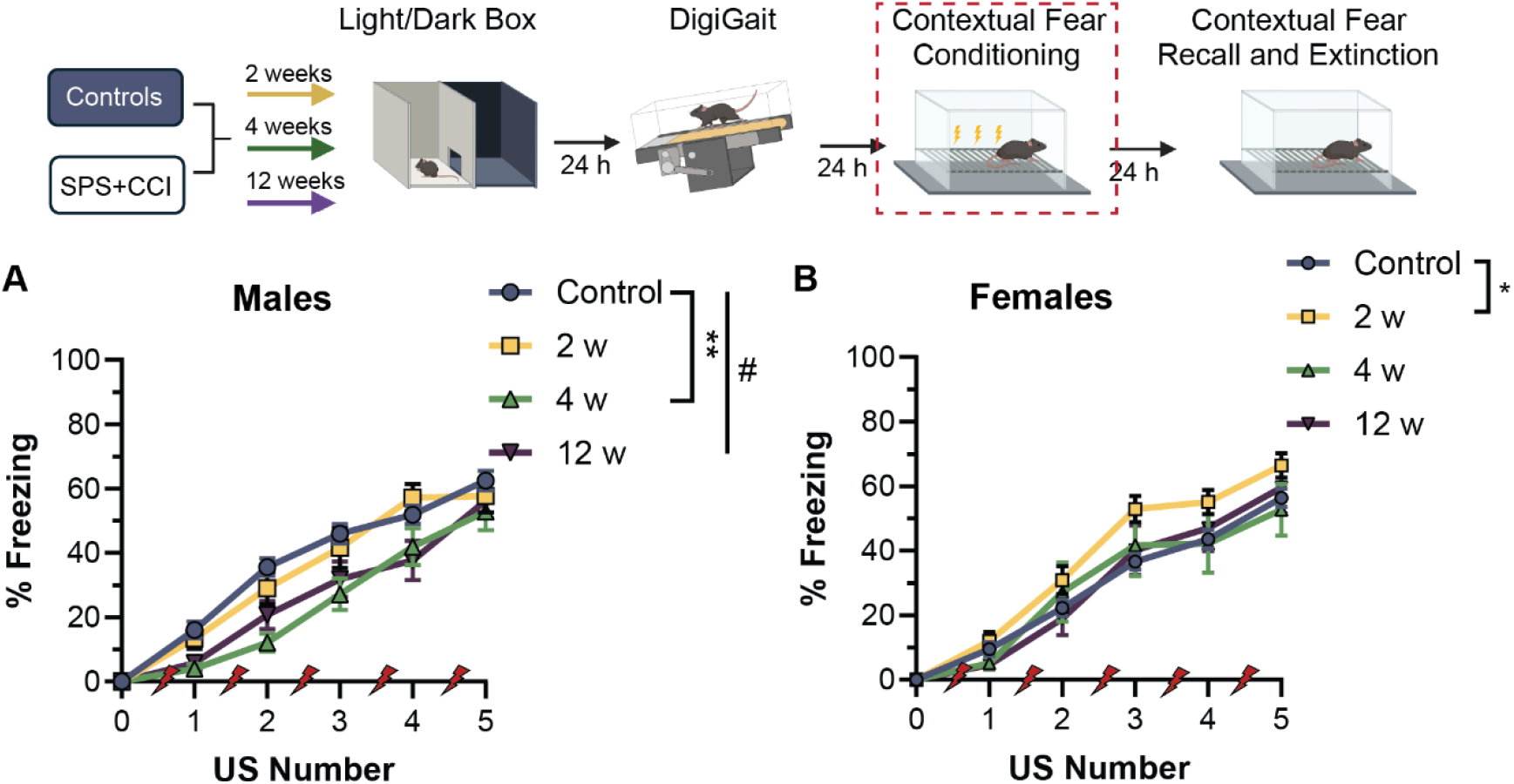
Delayed impairment of contextual fear acquisition in male mice after neurotrauma. A) There is a male specific learning deficit in contextual fear learning that emerges at 4 weeks and may persist until 12 weeks. B) Females acquire contextual fear faster 2 weeks post neurotrauma but normalize to Control levels by 4 weeks. All mice freeze at similar levels by the end of the fear conditioning session. **p<0.01, #p=0.09. Schematic prepared with Biorender.

Finally, we examined freezing at the end of contextual fear acquisition after delivery of the final foot shock (during the last 1 minute in the chamber), and found no significant differences between groups or sexes (between subjects, main effect of trauma group: *F*(3,143)=1.02, *p*=0.386; between subjects, main effect of sex: *F*(1,143)=0.15, *p*=0.702; trauma group x sex interaction: *F*(3,143)=1.16, *p*=0.326). This indicates that there were no trauma group or sex differences in final freezing levels or in the ability to express freezing behavior. The only difference observed was in the rate at which this freezing response was acquired.

### FC Day 2 - Extinction Freezing

24 hours after acquisition, mice were returned to the fear conditioning chamber for a 20-minute extinction session in the absence of any foot shock. Freezing was scored continuously and binned into 2-minute bins for analysis and visualization.

We first looked at freezing during the first 4 minutes of the extinction session on day 2 as an indicator of contextual fear recall using ANOVA. All neurotrauma groups froze significantly less than Controls (main effect of group: *F*(3,143)=25.74, *p*<0.001; post hoc Dunnett all *p* values <0.001 compared to Controls) and there were no effects of sex or interactions on fear recall (main effect of sex: *F*(1,143)=0.44, *p*=0.507; trauma group x sex interaction: *F*(3,143)=0.22, *p*=0.884) (**Fig. 5a**).

**Figure 5:**
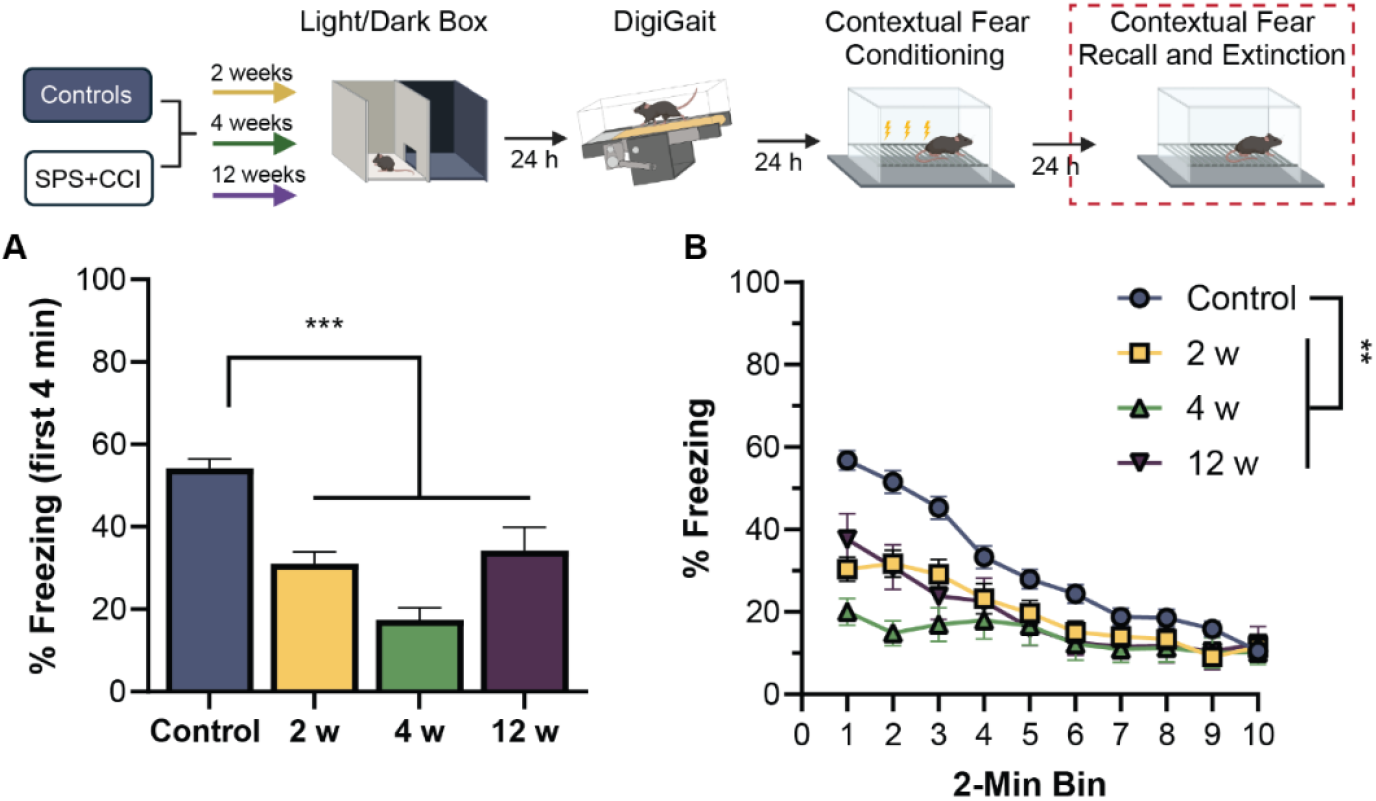
Impaired contextual fear recall in both sexes following neurotrauma. A) Contextual fear recall, quantified as percent freezing during the first 4 minutes of extinction, was significantly reduced in neurotrauma exposed mice at all time points relative to Controls (both sexes). B) Freezing during the entire extinction session in 2 minute bins revealed differences in extinction dynamics compared to Controls (both sexes). Data shown are mean +/-SEM. Asterisks indicate statistically significant differences. **p<0.01; ***p<0.001.

We then looked at freezing in 2-minute bins over the entire course of the extinction session using repeated measures ANOVA. There was a significant within subject effect of time during extinction (within subjects effect of time: *F*(5.42,774.42)=39.90, *p*<0.001) indicating that freezing changed over time while animals were exposed to the fear conditioning context. There was also a significant time x neurotrauma group interaction (within subjects, time x trauma interaction: *F*(16.25, 774.42)=5.84, *p*<0.001) with no within subjects effects of sex (time x sex interaction: *F*(5.42, 774.42)=0.95, *p*=0.454; time x sex x group interaction: *F*(16.25, 774.42)=0.53, *p*=0.943)(**Fig. 5b**). Overall, freezing during extinction in mice after neurotrauma was less than Controls (between subjects, main effect of neurotrauma: *F*(3,143)=10.93, *p*<0.001) with no effects of sex (between subjects, main effect of sex: *F*(1,143)=0.08, *p*=0.778; between subjects, sex x trauma group interaction: *F*(3,143)=0.18, *p*=0.913), consistent with earlier results describing impaired contextual fear recall. This impairment was present at all time points tested after neurotrauma compared to Controls (2 week: p=0.002; 4 week: p<0.001; 12 week: p=0.008, Dunnet’s).

To more closely examine learning rates during the extinction session we used GraphPad Prism to generate nonlinear regression curve fits (robust) to each animal’s extinction data to produce a slope value for each animal (**Fig 6A**). Due to differences in freezing at the beginning of the extinction session we used normalized freezing ratios calculated by dividing the freezing in each 2-minute bin by each animal’s average freezing in the first 4 minutes of extinction (**Fig. 6B, 6C**). This allows us to control for existing differences in memory recall and makes slopes comparable across animals. These slopes were then analyzed using ANOVAs (between subjects factors = neurotrauma group, sex). Although we did not find a significant main effect of neurotrauma timepoint (between subjects, main effect of group: *F*(3,142)=1.37, *p*=0.256) there was a significant sex x trauma group interaction (sex x trauma interaction: *F*(3,142)=4.81, *p*=0.003) and a significant main effect of sex on extinction slope (main effect of sex: *F*(1,142)=9.03, *p*=0.003). Due to the significant interaction, we did follow up ANOVAs on each sex separately. We found no effect of neurotrauma on extinction slope in female mice (main effect of group: *F*(3,65)=0.62, *p*=0.603). In males, however, there was a significant effect of group on extinction slope (main effect of group: *F*(3,81)=6.10, *p*<0.001) that Dunnett’s tests revealed was driven by flatter extinction curves in the 4 week time point (*p*<0.001). Neither the 2 week timepoints nor the 12 week timepoints revealed any significant effects on extinction slope compared to Controls (2 week: *p*=0.292; 12 week: *p*=0.212, Dunnet) (**Fig. 6A**).

**Figure 6:**
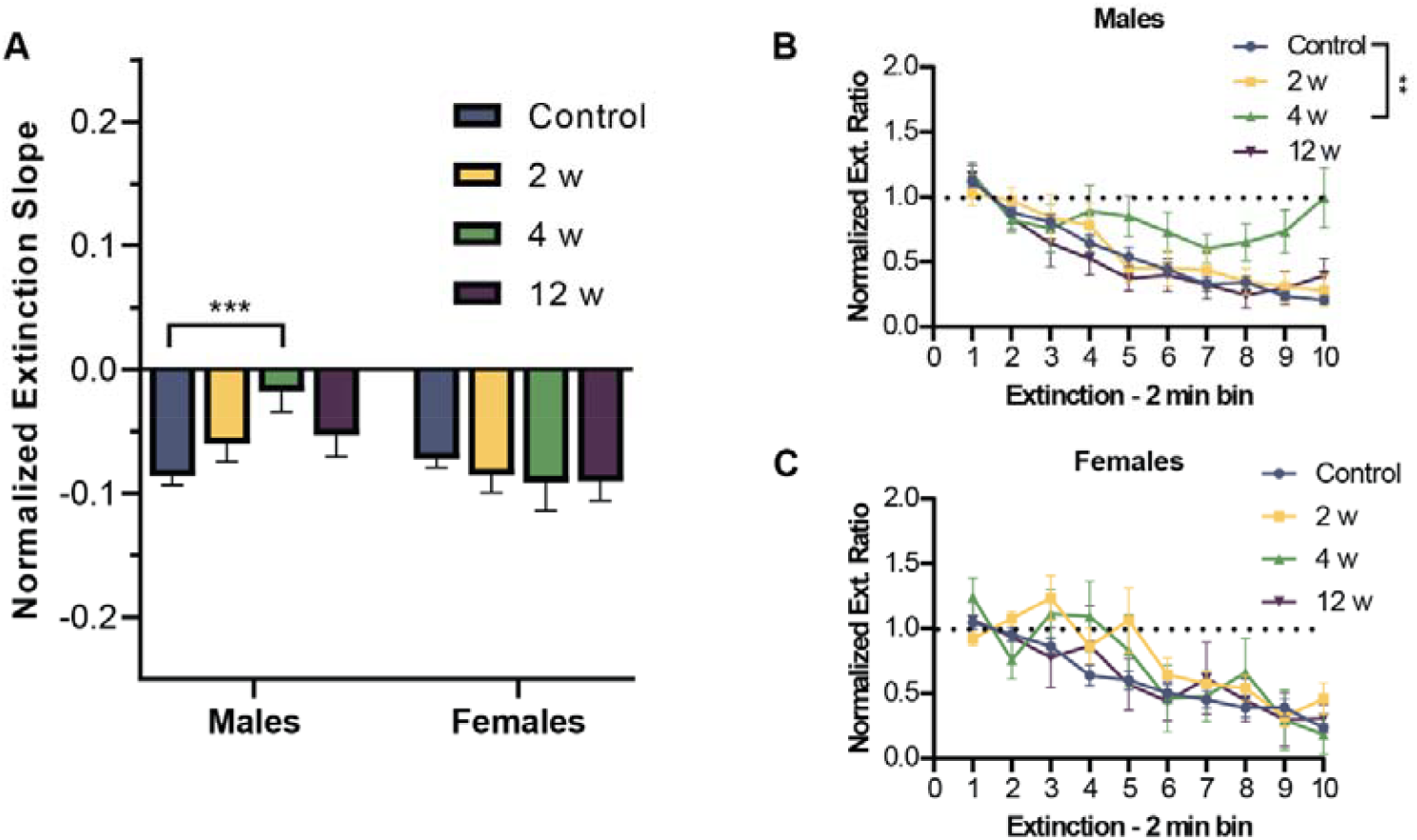
Impaired extinction rate in males emerges at 4 weeks. A) Normalized freezing for each mouse was analyzed by calculating a freezing ratio for each bin by dividing by their freezing in the first 4 min of extinction. B,C) Males but not females showed flatter extinction slopes at 4 weeks post neurotrauma. Dotted line indicates no change in freezing during extinction session (compared to first 4 minutes).

### Behavioral Predictors of Cognition

We then wanted to explore whether any of these behavioral phenotypes predicted cognitive ability in animals exposed to neurotrauma. Exploratory linear regressions (Enter method) were performed on each of the cognitive outcomes described here: contextual fear recall (freezing during the first 4 minutes of extinction) and learning rate (normalized extinction slope) with all of the variables reported here (dark duration in the LD box, number of light entries in the LD box, fore and hind stance width during DigiGait, and left and right paw placement positioning during DigiGait) entered into the model. Due to the exploratory nature of this analysis no variables were removed from the regression model. Due to the relatively small sample size of subjects that completed all tests with useable data all time points and both sexes were included. None of the behavioral variables significantly accounted for variance in contextual fear recall (overall ANOVA: *F*(6,31)=0.33, *p*=0.916). However, the regression model was significant in predicting extinction slope (overall ANOVA: *F*(6,30)=3.69, *p*=0.007, adjusted R^2^=0.309). Paw placement positioning for both paws during DigiGait were significant predictors of extinction slope (Left: B=-0.197, *t*=-3.040, *p*=0.005; Right: B=-0.120, *t*=-2.057, *p*=0.048), whereas no other measures significantly predicted extinction slope (all *p* values >0.21). Follow up simple linear regressions on each timepoint for each sex were conducted for left PPP and extinction slope. We found that this effect was driven by males in the 4 week (R^2^=0.6512, *p*=0.0155) and 12 week (R^2^=0.7525, *p*=0.0114) timepoints (2 week: R^2^=0.3546, *p*=0.1583) and not females (all *p* values >0.3), consistent with the emergence of the extinction deficit in these groups (Fig 7. A,B).

**Figure 7:**
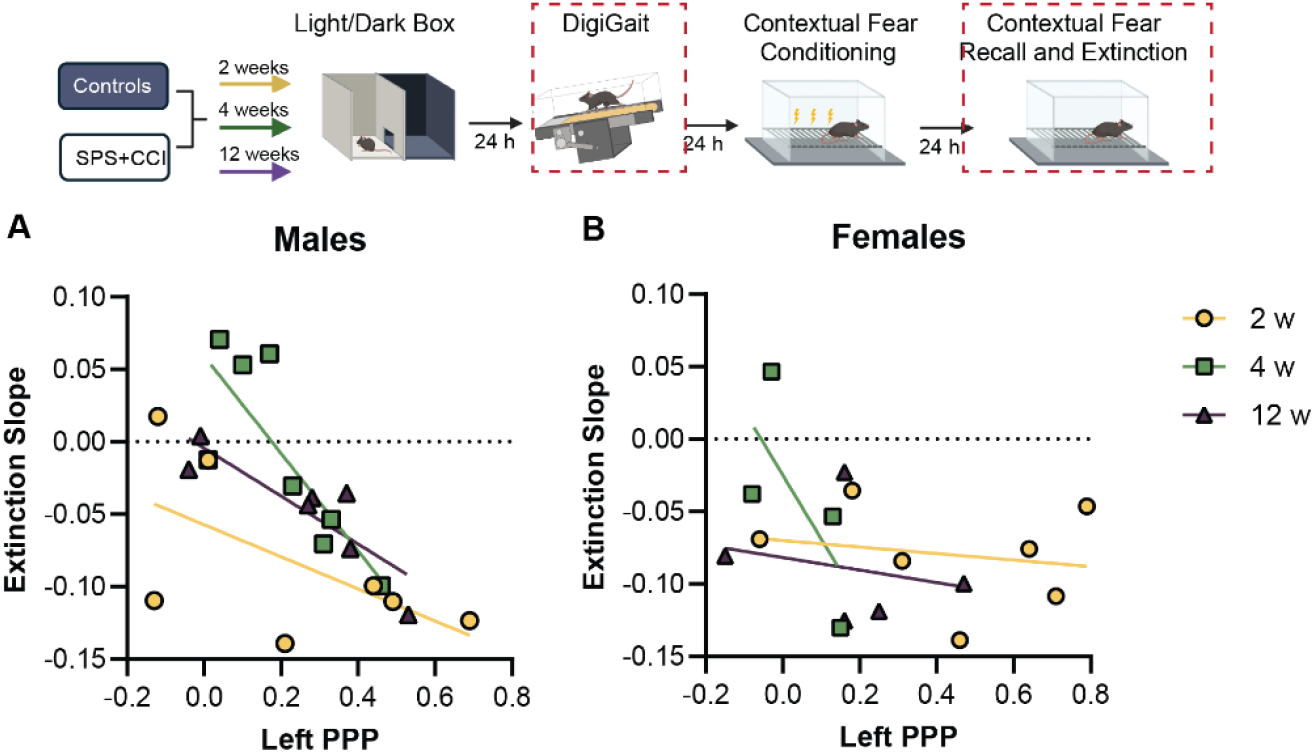
Relationship between gait (contralateral paw placement positioning) and cognition (normalized extinction slope) after combined SPS+CCI. A) Starting 4 weeks post-SPS+CCI, males with lower PPP values, indicative of balance impairment, also had higher extinction slope values, indicative of impaired learning. B) There was no relationship between PPP and extinction slope in females at any time point.

## Discussion

The present findings demonstrate that our model of combined neurotrauma (SPS+CCI) produced sex and time dependent behavioral deficits in mice. Our results found that combined neurotrauma produced early and persistent gait impairments in both sexes emerging at 2 weeks post-injury, delayed changes to anxiety-like behavior characterized by reduced avoidance of an anxiogenic environment emerging at 12 weeks post-injury, and long-lasting contextual fear recall deficits at 2, 4, and 12 weeks post-injury. Males, but not females, showed impaired learning, with reduced rates of fear acquisition and extinction emerging at 4 weeks post-injury, consistent with previous clinical and preclinical studies showing sex differences in neurobiological responses to trauma.^33, 51-53^ This work highlights the complexity and necessity of modeling ecologically valid disease conditions such as comorbid PTSD+TBI in human military populations and lays an important benchmark against which potential treatment outcomes could be measured in this mouse model.

### Balance and Gait Impairments Emerge Early and Persist

Measures of gait, including paw placement positioning (PPP) and stance width, were used as objective outcomes of balance and postural stability. In our model, gait deficits appeared early and persisted across all time points. PPP reflects the overlap between ipsilateral fore and hind limb placement. Higher stance width in the fore limbs paired with lower PPP suggest unstable gait and compensatory bracing as the mice run downhill. These results are consistent with previous work examining gait metrics after TBI.^45, 54, 55^

Clinical studies report that individuals with neurotrauma frequently experience balance problems and motor deficits, particularly under dual-task conditions.^56-58^ Our observation that male mice with slower fear extinction also had worse PPP suggests that early balance impairments may reflect vulnerability of neural systems involved in both motor and cognitive control and support a relationship between gait and cognition.^59-62^ This is consistent with the observation that subtle gait or balance abnormalities emerge early, during prodromal phases, of neurodegenerative diseases such as Parkinson’s disease.^63-67^ Therefore, PPP may represent a translational behavioral marker, analogous to the early subtle gait disturbances in humans, and providing a useful marker of prodromal neurodegenerative vulnerability as well as a measurable end point for evaluating therapeutic interventions in rodent models of combined neurotrauma.

### Delayed Emergence of Blunted Emotional Reactivity in the LD Box

At 12 weeks post-neurotrauma, mice did not show the typical preference for the dark region of the Light/Dark box. While increased time in the light zone is conventionally interpreted as reduced anxiety, TBI literature suggests that blunted emotional reactivity and impaired threat processing can also drive such outcomes,^68, 69^ making mice less responsive to anxiogenic cues (like bright light), not necessarily less anxious. Unlike acute anxiolytic phenotypes, chronic effects of TBI can disrupt sensory and threat circuits (e.g. amygdala and hippocampus), resulting in diminished responding to anxiogenic stimuli.^70, 71^

Rodent TBI studies show mixed anxiety-like profiles over time, with some reporting early increases in anxiety-like behavior, followed by a shift towards increased exploration in anxiogenic zones at later post-injury intervals (>5-7 weeks), supporting the idea of evolving neural dysfunction.^70, 71^ Importantly, in the current study, we found no differences in the number of light entries, suggesting that neither hyperactivity nor impulsivity is likely to underlie this change. This interpretation is supported by clinical observations that patients with neurotrauma may exhibit reduced emotional responsiveness and altered risk evaluation, which can manifest as poor threat discrimination or indifference.^72-74^

Of note, the literature is not uniformly consistent regarding PTSD models and anxiety-like behaviors, particularly in SPS models, which may not induce increased anxiety like behavior as expected.^75, 76^ Alternatively, the lack of any side preference, paired with the cognitive deficits we observed in fear extinction, suggests that these mice may not meaningfully interpret the contextual cues of the light and dark chambers during the chronic stages of neurotrauma recovery.

### Cognitive Dysfunction: Learning, Recall, and Sex Differences

The underlying neural circuitry of contextual fear conditioning and extinction (including the hippocampus, PFC, and amygdala^77, 78^) are known to be vulnerable to both physical and psychological trauma with prior work demonstrating that disruptions in these regions after either TBI or PTSD can impair memory retrieval and the extinction process.^79-83^ Our data reveal two separable components of cognitive impairment after neurotrauma: learning deficits that emerge at 4 weeks in males in both initial learning and extinction, and memory recall deficits evident at all time points in both sexes.

Fear acquisition and recall: male mice exposed to SPS+CCI displayed slower contextual fear acquisition relative to sex matched Controls by 4 weeks, with a trend towards persistence at 12 weeks. Female mice did not exhibit this learning deficit; instead, they showed transiently heightened freezing to foot shocks at 2 weeks post SPS+CCI, whereas freezing levels at 4 weeks post SPS+CCI were comparable to Controls. All mice eventually reached equal levels of freezing during the initial fear acquisition session, indicating that 1) freezing is physically possible for these animals and 2) despite slower learning rates, all animals reach the same final level of freezing, indicative of intact learning ability. However, both males and females exhibited impaired contextual fear recall at each tested interval after neurotrauma, echoing findings from our previous work demonstrating that this combined SPS+CCI model disrupts recall of contextual fear memory.^45^

Extinction: Despite lower initial levels of freezing during extinction, male mice exposed to SPS+CCI displayed slower rates of contextual fear extinction relative to sex matched Controls. Similar to learning rates during acquisition, this learning impairment was strongest at 4 weeks but still trending at 12 weeks post SPS+CCI. Sex differences in fear extinction and its modulation by gonadal hormones are well documented in rodents and humans,^84-87^ with implications for how males and females engage overlapping but distinct neural circuits during fear learning and extinction.^88, 89^ Fear extinction deficits are also a feature of PTSD in clinical settings.^80, 83^ In particular, SPS has been shown to disrupt extinction across contextual and cued fear without necessarily altering initial acquisition^5, 47, 90^ with effects more pronounced in males.^13, 91^

### Sex Specific Relationship Between Motor and Cognitive Outcomes

Beginning at 4 weeks post-injury, male mice with lower contralateral paw placement positioning (PPP), an indicator of greater balance impairment, also exhibited flatter extinction slopes, an indicator of slower extinction learning. This coupling between gait instability and impaired fear extinction was not observed at earlier time points and was absent in females at all intervals examined. The relationship between gait and cognitive function after neurotrauma observed in males aligns with clinical evidence in humans that motor and cognitive deficits frequently co-occur and are interrelated following brain injury. In adults recovering from TBI, measures of gait and complex motor performance have been shown to correlate with cognitive performance, particularly in tasks that place concurrent demands on attention, executive control, and motor coordination.^92, 93^

Other studies have further found correlations in dual task gait performance and cognitive decline in men but not women supporting the idea of a sex moderation of the gait-cognition relationship.^94, 95^ Our observation of a meaningful individual difference correlation in this mouse model not only provides insight into how combined neurotrauma affects underlying neural systems, but also enhances the ecological validity of the model relative to the human condition, where the relationship between motor and cognitive deficits in neurological disease is well established.^96-99^

### Potential Mechanisms

Our behavioral results are consistent with previous work on PTSD-relevant paradigms that disrupt hippocampal-prefrontal-amygdala circuitry critical for contextual fear learning and extinction.^14, 100^ Combined psychological stress and physical brain injury may exacerbate dysfunction within these corticolimbic networks that integrate emotional and motor behavior resulting in comorbid gait and cognitive impairment. The emergence of maximal impairment at 4 weeks post-neurotrauma may be driven by secondary processes including chronic neuroinflammation, synaptic remodeling, and/or altered neuromodulatory signaling, that progressively develop over the weeks following injury.^33, 101^ Our results are consistent with the gradual worsening of both PTSD and TBI behavioral symptoms in some populations (particularly military personnel) of human patients exposed to trauma.^24, 102-104^ Preclinical studies further indicate that sex differences in behavioral outcomes and neuroinflammatory responses after experimental TBI are common, with males often showing greater vulnerability to chronic dysfunction, although findings vary across models and outcome measures.^33, 101^

### Considerations and Future Work

Limitations of this study include several considerations. First, fear based cognitive assays are inherently aversive and may inadvertently model aspects of PTSD re-exposure.^105, 106^ However, many traditional cognitive tests for rodents require precise movements or motor coordination, which confound interpretation in subjects with balance or gait impairments. Careful design and validation of non-aversive cognitive assays that minimize motor demand will be an important direction for future work. Second, as with most preclinical models, experimental manipulations in mice are highly controlled and cannot capture the full spectrum of biological, genetic, or experience-based heterogeneity that characterizes comorbid PTSD and TBI in humans. Nonetheless, despite this controlled setting, we observed substantial inter-individual variability across behavioral domains and time points, suggesting that meaningful differences in vulnerability and recovery trajectories emerge even under these standardized conditions. Such variability may be particularly informative for identifying phenotypes relevant to treatment response.

Future work will examine neurobiological markers of neurodegeneration and circuit dysfunction to bridge behavioral phenotypes with underlying neuropathology and continue to examine both sexes to better define sensitive markers for females.

Finally, this study was not designed to dissociate the independent effects of PTSD and TBI, but rather to characterize the temporal and sex-dependent behavioral profile that emerges with the comorbid condition. Given the high prevalence of comorbid PTSD and TBI in clinical populations, this approach prioritizes ecological validity and establishes a foundation for systematic testing of intervention timing and efficacy in a preclinical setting.

## Conclusion

The present findings demonstrate sex- and time-dependent impairments in gait and contextual fear learning following combined PTSD and TBI in mice, with peak deficits emerging at 4 weeks post-neurotrauma. Male mice exhibited impaired contextual fear acquisition and extinction, whereas both sexes showed deficits in contextual fear recall and gait. These results extend our prior work using the SPS+CCI model^45^ by defining the temporal progression and sex specificity of behavioral dysfunction following combined neurotrauma.

By identifying reproducible motor and cognitive phenotypes across defined post-injury intervals in both sexes, this mouse model provides a foundation for higher-throughput screening of candidate interventions and for testing treatment timing and sex-specific efficacy in a manner that more closely reflects the clinical reality. Given the high prevalence of comorbid TBI and PTSD in clinical populations, particularly in military and Veteran cohorts,^107, 108^ and epidemiological evidence linking neurotrauma to increased risk of Parkinson’s disease and related synucleinopathies ^37-39, 109-112^ experimental models that capture the interaction between physical brain injury and psychological trauma are well positioned to inform understanding of long-term functional trajectories and identify windows of risk or treatment opportunity following neurotrauma.

